# Epigenetic bases of grafting-induced vigour in eggplant

**DOI:** 10.1101/831719

**Authors:** Elisa Cerruti, Carmina Gisbert, Hajk-Georg Drost, Danila Valentino, Ezio Portis, Lorenzo Barchi, Jaime Prohens, Sergio Lanteri, Cinzia Comino, Marco Catoni

## Abstract

In horticulture, grafting is a popular technique used to combine positive traits from two different plants. This is achieved by joining the plant top part (scion) onto a rootstock which contains the stem and roots. Despite its wide use, the biological mechanisms driving rootstock-induced alterations of the scion phenotype remain largely unknown. Given that epigenetics plays a crucial role during distance signalling in plants, we studied the genome-wide changes induced by DNA methylation in eggplant (*Solanum melongena*) plants grafted onto two interspecific rootstocks used to increase scion vigour. As a control, we compared any epigenetic effect found in such grafts to patterns detected in self-grafted plants. We found that vigour was associated with a specific change in scion gene expression and a genome-wide hypomethylation in CHH context. Interestingly, this hypomethylation correlated with the down-regulation of younger and potentially more active LTR retrotransposons (LTR-TEs), suggesting that graft-induced epigenetic modifications are associated to both physiological and molecular phenotypes in grafted plants. We propose that rootstocks can promote vigour by reducing DNA methylation in the scion genome, following similar principles found in some heterotic hybrids.

## Introduction

Grafting is the process of joining plant tissues of two plants: the scion (upper part) and rootstock (lower part), which then continue to grow together combining the favourable characteristics of the genotypes involved. Plant grafting is a naturally occurring process, but was systematically used by humans as an agricultural technique for fruit plant trees. Over centuries, grafting allowed humans to facilitate tree propagation, reduce juvenility, provide resistance to biotic and abiotic stresses, and to control plant growth (Gautier *et al.*, 2018). Starting from the early twentieth century grafting has been also extensively used in vegetable, mainly in Solanaceae and Cucurbitaceae species, making also use of interspecific combinations (Goldschmidt, 2014). Although not directly involved in fruit production, rootstocks are selected for their ability to regulate salinity and drought tolerance, water-use efficiency and nutrient uptake, soil-borne pathogen resistance, scion vigour and architecture, mineral element composition, fruit quality and yield in a broad range of species (Colla et al., 2017; Kumar et al., 2017; Warschefsky et al., 2016). Despite its agricultural success in the past centuries, the molecular mechanisms at the base of rootstock-mediated control of scions phenotype remain mostly unknown.

So far, several studies on model species demonstrated that a possible molecular mechanism involved in grafting is a bi-directional long-distance transport of mRNAs transcripts and signalling macromolecules such as microRNAs (miRNAs) and small RNAs (sRNAs). These RNAs are able to trigger physiological changes through the graft junction (Bai et al., 2011; Lewsey et al., 2016; Melnyk et al., 2011; Molnar et al., 2010; Thieme et al., 2015) which might also result in emergence of vigour. Using grafting as experimental system, the role of epigenetics in this process was suggested after the finding that sRNAs are able to induce epigenetic variation through the RNA directed DNA methylation (RdDM) pathway in recipient tissues (Lewsey et al., 2016; Melnyk et al., 2011; Zhang et al., 2018), providing the first molecular basis for grafting-induced changes in organ growth and development. However, a defined molecular mechanism driving rootstock-dependent physiological effects in grafted plants still remains elusive.

Here, we introduce the eggplant (*Solanum melongena* L., 2n = 2x = 24) as a model system to study general molecular mechanisms of grafting. Eggplant, is one of the most common commercially grafted solanaceous crops (Lee and Oda, 2010). Several rootstocks were selected to improve the quality of eggplant cultivation, providing resistance/tolerance to soil pathogens and inducing vigorous growth of the scions (Gisbert et al., 2011; Schwarz et al., 2010). Commercial grafting of eggplants also makes large use of interspecific combinations to increase plant vigour and resistance to pathogens, and the most popular rootstocks include wild solanaceous species, like the eggplant wild relative *Solanum torvum* (Turkey berry), or tomato hybrids developed specifically for being used as rootstock (Bogoescu and Doltu, 2015; Miceli et al., 2014). A recent study provided the first evidence of locus-specific changes in DNA methylation in inter-species grafting of *S. melongena* and other Solanaceaes (Wu et al., 2013), suggesting that rootstock-induced epigenetic alterations can produce physiological changes in eggplant scions.

Here we analysed the genome-wide methylation profiles of eggplant scions from interspecific grafting combinations, using *S. torvum* and a tomato hybrid as rootstocks. We observed that the enhanced vigour induced by these rootstocks is associated with genome-wide decrease of CHH methylation, occurring at both coding genes and transposons. In addition, we found that DNA demethylation is also associated with a difference in transcriptional analysis between hetero-grafted and self-grafted plants. Interestingly, many of the identified differentially regulated genes are involved in plant developmental processes directly or indirectly related to the grafting response while Transposable Elements (TEs) transcriptional regulation appears to be modulated in an age-related fashion.

## Results

### Enhanced vigour in hetero-grafted eggplant scions is associated to genome-wide CHH hypomethylation

To study the effect of grafting on vigour, we grafted eggplant scions (double haploid line derived from the commercial hybrid ‘Ecavi’) on three rootstocks: i) the wild species *Solanum torvum*, ii) the tomato F1 commercial hybrid ‘Emperador RZ’ and iii) the same eggplant genotype (self-grafting) (**Fig. 1a**). Both *S. torvum* and ‘Emperador RZ’ were previously reported to induce vigour in eggplant scions (Bogoescu and Doltu, 2015; Gisbert et al., 2011). Indeed, five months after grafting, the hetero-grafted plants showed a remarkable and statistically significant increase in height when compared to self-grafted eggplants, used as reference control (**Fig. 1b**). Depending on whether the rootstock used was *S. torvum* or the tomato ‘Emperador RZ’, eggplant scions respectively displayed a marked bushy phenotype or more pronounced vertical growth (**Fig. 1c**).

**Figure 1.**
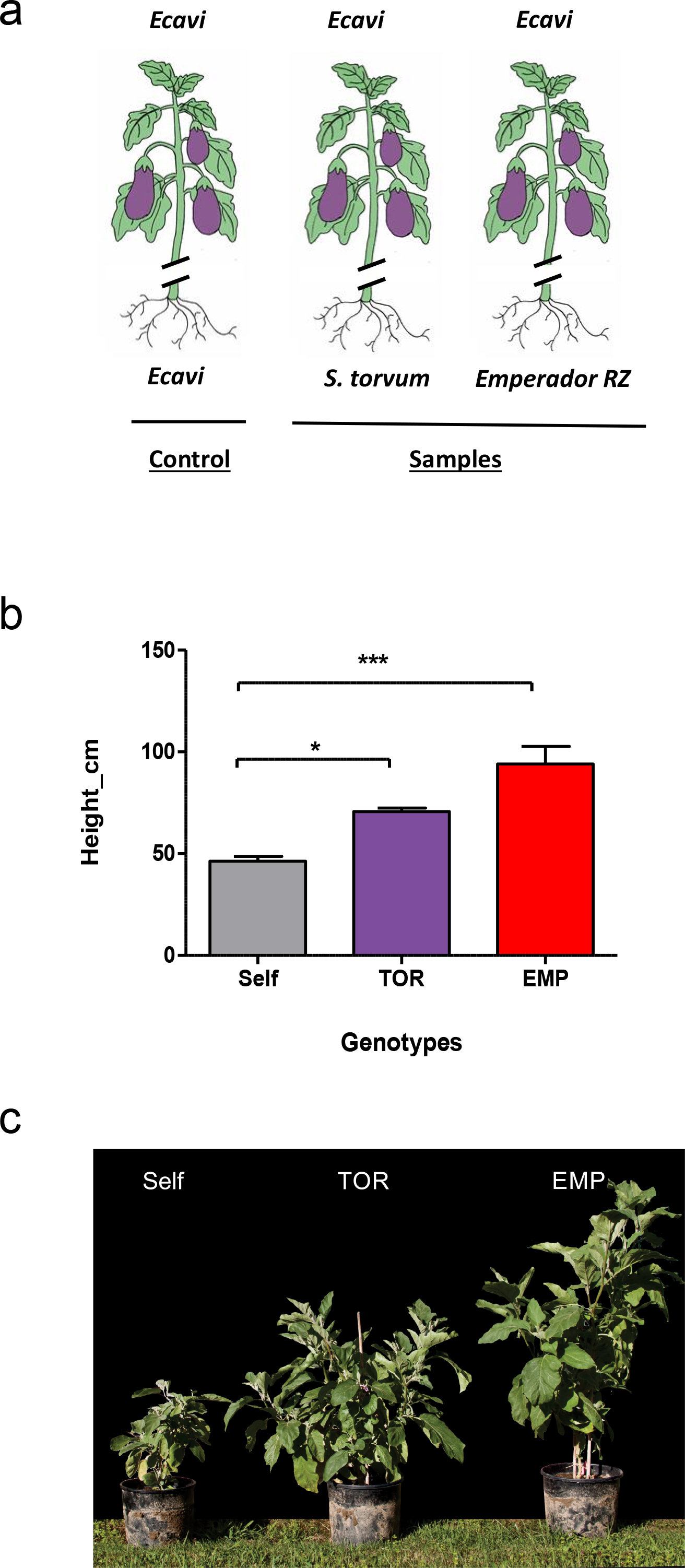
Heterografting induces vigour in eggplant scions. **a** Representation of grafting experimental design considering the conditions under investigation (from left to right): 1) self-grafted eggplant (var. ‘Ecavi’) scions, 2) ‘Ecavi’ scions onto *S. torvum* rootstocks (TOR) and 3) ‘Ecavi’ scions onto tomato ‘Emperador RZ’ rootstocks (EMP). **b** Height differences between hetero-grafted and self-grafted plants observed five months after grafting occurred. From left to right eggplant scions grafted on ‘Ecavi’ eggplant (Self), *S. torvum*’ (TOR), and tomato ‘Emperador RZ’ (EMP). Asterisks mark statistically significant differences (ANOVA 1-way, *P* < 0,05). Error bars represent SD of three replicates. **c** Picture displaying differences in vigour between plants representative of the grafting conditions at five months post grafting.

It has been hypothesised that DNA methylation is the driving mechanism generating phenotypic diversity via grafting (Melnyk, 2017). To test whether changes in cytosine methylation might indeed be associated with observed differences in scion vigour, we performed genome-wide bisulfite sequencing of DNA samples extracted from two eggplant scions of each grafting combination described above, and two biological replicates of ungrafted eggplants. After quality control and data processing, roughly 72.5 M reads per sample were sequenced, with an average 9× coverage of the eggplant genome. We generated the first eggplant genome methylation profile at single cytosine resolution, which in leaf tissue displayed 91% methylation in CG, 84% in CHG and 19% in CHH contexts (**Table S1**). In wild type ungrafted eggplant, the DNA methylation in CG and CHG contexts was more pronounced in the central part of each chromosome, while decreased occurred in the terminal parts of the chromosome arms. In contrast, CHH methylation was more evenly distributed across the genome (**Fig. S1**). This profile is similar to DNA methylation patterns reported for the same tissue in other Solanaceaes (Wang et al., 2018; Zhong et al., 2013), characterized by a general anti-correlation of DNA methylation (mostly in CG and CHG context) and coding genes and is associated with an increasing abundance of methylated TEs in the central part of chromosomes (**Fig. S1**).

While the methylation profile of self-grafted scions was similar to ungrafted plants (**Table S1**), when we checked methylation profiles in hetero-grafted scions we observed a significant genome-wide decrease in CHH methylation of 3.37% and 2.58% respectively in scions grafted onto *S. torvum* and ‘Emperador RZ’ when compared to the self-grafted plants (**Fig. 2a**). This decrease appeared to be uniformly distributed along chromosomes (**Fig 2b**, **Fig. S2**). Unlike methylation in CHH context, the methylation in CG and CHG contexts remained unchanged in both self- and hetero-grafted scions (**Fig. 2a-b, Fig. S2**). Further analyses showed that CHH hypomethylation was more prominent at TEs than at coding genes, but not specific for a particular TE family (**Fig. 2c-d**, **Fig. S3**).

**Figure 2.**
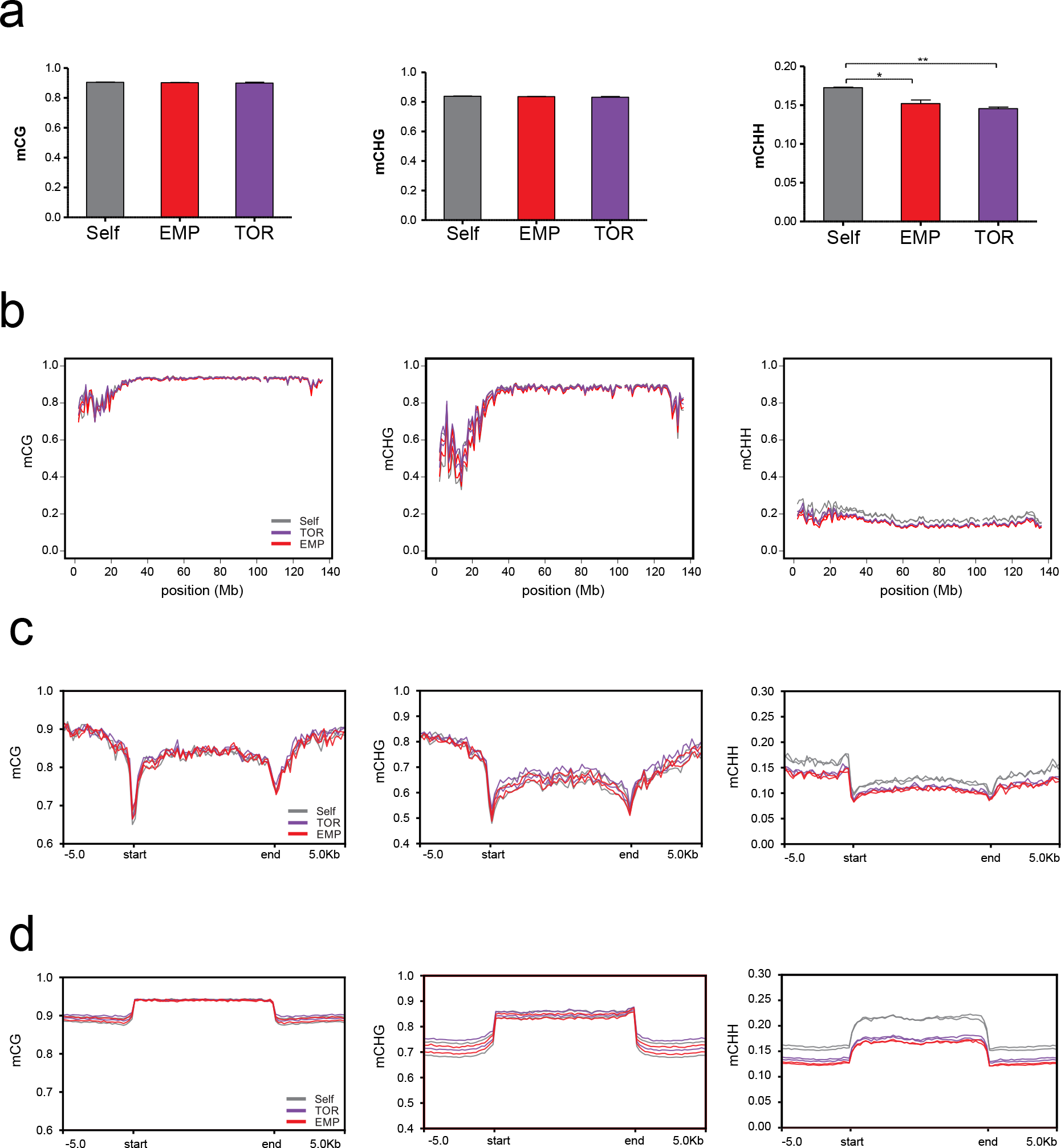
Heterografting is associated to CHH genome hypomethylation. **a** Methylation averaged at all cytosines for two replicates per hetero-grafted condition. Error bars represent SD (*n*=2). * = t-test p-value * = 0.0005, ** = 0.0007. **b** Distribution of DNA methylation at the three cytosine contexts (mCG, mCHG and mCHH) along chromosome 1 of eggplant genome in eggplant scions self-grafted (Self), grafted on *S. torvum* (TOR) or grafted on tomato ‘Emperador RZ’ (EMP). Information at the remaining chromosomes (2 to 12) are displayed in Figure S2. **c** Distribution of averaged DNA methylation at annotated genes in the three contexts (mCG, mCHG and mCHH,) in the eggplant scions self-grafted (Self), grafted on *S. torvum* (TOR) or grafted on tomato ‘Emperador RZ’ (EMP). **d** Distribution of averaged DNA methylation level at annotated repetitive elements, description is as in c).

### Hetero-grafted plants display similar transcriptional profile

To further investigate whether differences in DNA methylation were associated with changes in transcription, we profiled the genome-wide RNA expression in the same grafted scion samples, by strand-specific RNA-sequencing (**Table S2**). Next, we compared the transcriptome of both eggplant hetero-grafted scions to self-grafted scions. Despite different species were used as rootstock, we observed that the transcription profiles of eggplant scions grafted onto *S. torvum* and ‘Emperador RZ’ clustered together and clearly diverge from self-grafted controls (**Fig. 3a**). This indicates that both changes in DNA methylation and in transcription are associated with the altered phenotype observed in hetero-grafted eggplants compared to self-grafted plants. The differential expression analysis revealed a prevalence of down-regulated genes in scions grafted onto both *S. torvum* (65%, **Fig. 3b**) and ‘Emperador RZ’ (61%, **Fig 3c**). In particular, we observed that 464 genes were up-regulated and 875 were down-regulated in scions grafted onto *S. torvum*, while 434 and 704 genes were found respectively up and down-regulated in scions grafted onto ‘Emperador RZ’ (**Fig. S4 a-b**, **Tables S3-S4**). In addition, 151 up-regulated and 462 down-regulated genes are shared between the two hetero-grafted categories (**Fig. S4 a-b**, **Tables S3-S4**). Validation performed by qPCR confirmed the change of expressions of 11 randomly selected genes, in scions grafted onto *S. torvum* (6 genes) and onto ‘Emperador RZ’ (5 genes) (**Fig. S5, Table S5**).

**Figure 3.**
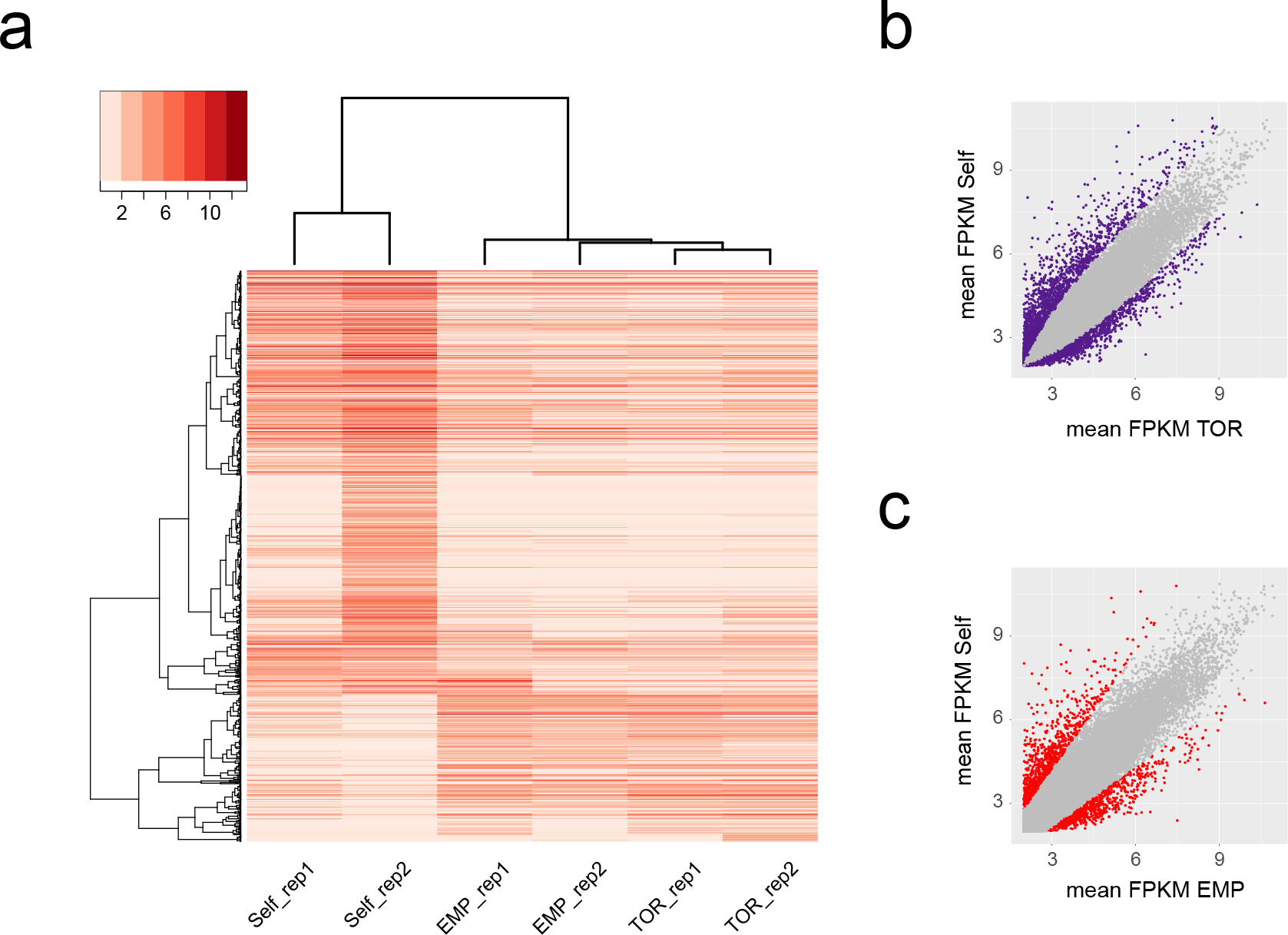
Hetero-grafted scions display similar gene expression profiles. **a** Heatmap illustrating the expression (log2 FPKM) of differentially expressed genes (log2 > 2 and FDR <0.05) in hetero-grafted plants and self-grafted controls. Each row represents one gene (*n* = 773). Coloured bars on the top-left indicate the expression level. Each column represents an eggplant scion grafted on different rootstock (TOR = *S. torvum*, EMP = tomato ‘Emperador RZ’ and Self = self-grafted). Expression values were ordered according to hierarchical clustering (hclust and heatmap3 in R software environment). The Euclidean distance dendrogram is presented on the left. **b** Scatter-plots of annotated genes (*n* =34,917) in Self and EMP conditions, each dot represent a genes and in red are displayed differentially expressed genes between the two conditions. **c** Scatter-plots of annotated genes in Self and TOR conditions. In purple are displayed differentially expressed genes between the two conditions.

Next, we explored whether the decrease in CHH methylation observed in the two hetero-grafted conditions is associated with a change in expression level of key regulators of non-CG methylation pathways. For this purpose, we selected a list of the most relevant genes involved in RdDM processes in *Arabidopsis* and, using a *blastp* search, we retrieved the corresponding eggplant orthologs (**Table S6**) and assessed their transcriptional status. We observed that the eggplant orthologs of the nucleosome remodeler DDM1 (SMEL_002g159290.1) and the argonaute protein AGO7 (SMEL_001g147190.1) were up-regulated in both eggplant grafted on *S. torvum* and ‘Emperador RZ’, compared to the self-grafted controls (**Table S6**), suggesting that epigenetic regulation could be differentially modulated in hetero-grafted plants.

In order to examine whether differentially expressed genes (DEGs) were involved in specific developmental processes that could explain the vigour of hetero-grafted scions, we performed a GO-enrichment analysis taking into account, separately, two datasets containing respectively up- and down-regulated genes in eggplants grafted on *S. torvum* and ‘Emperador RZ’ using the ShinyGO tool (Ge *et al*, 2018; **Tables S7-S8**). We observed enrichment (p-value < 0.05) among up-regulated genes involved in early developmental processes such as cell division, regulation of cell-cycle and DNA replication, which are consistent with the increased vigour observed in the hetero-grafted plants (**Table S7, Fig S6**). Side by side, the enrichment analysis on down-regulated genes (p-value < 0.05) highlighted a basal response characterised by prevalent GO terms associated to transmembrane transport, ion binding and response to stimuli, which might be directly or indirectly triggered by grafting (**Table S8, Fig S6**).

### Grafting modulates Transposable Elements expression

We then inspected whether transcriptional activity of TEs, which are directly silenced by DNA methylation, differed between hetero-grafted and self-grafted plants. After filtering repeats annotated on the eggplant reference genome (Barchi et al., 2019), we selected putative transposable elements (**Table S9**) and performed differential expression analysis. TEs appeared to be regulated similarly to genes, with more down-regulated elements (341 and 201 TEs down-regulated and 95 and 32 TEs up-regulated in scions grafted respectively onto *S. torvum* and ‘Emperador RZ’) compared to the self-grafted plants (**Table S10**). Specifically, we observed that non-LTR retrotransposons belonging to the RTE class were the most abundant among up-regulated TEs in scions grafted both onto *S. torvum* (53 TEs) and ‘Emperador RZ’ (16 TEs). On the other hand, the most represented down-regulated TEs were LTR retrotransposon of the Gypsy superfamily, (106 and 119 TEs in ‘Emperador RZ’ and *S. torvum* hetero-grafted plants respectively) of which, a significant proportion (105 TEs) were commonly repressed in both hetero-grafted scions **(Fig. 4a-b, Table S10)**.

**Figure 4.**
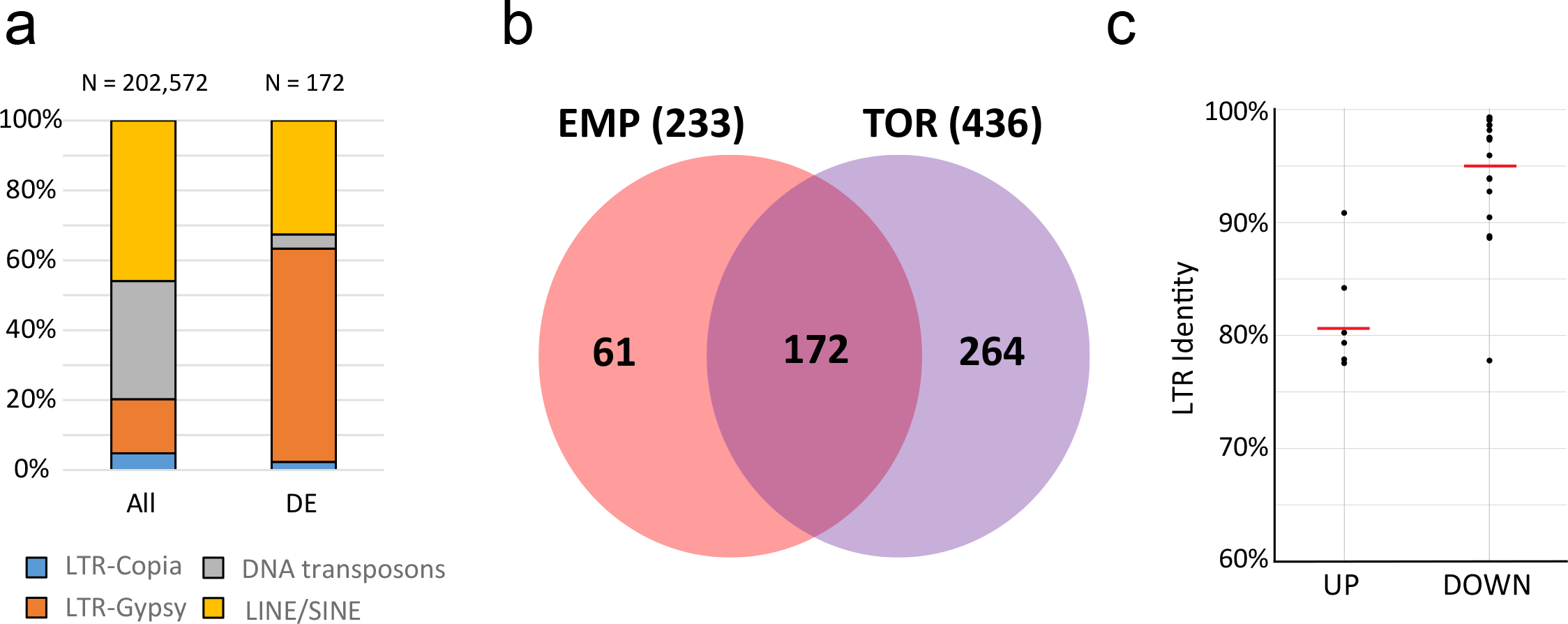
LTR-TEs regulation during heterografting. **a** Bar plot displays the distribution of families of TE differentially expressed (DE) in hetero-grafted combinations (compared to self-grafted controls). The composition of TEs families for all annotated elements is reported as comparison (All). **b** Venn-diagram displaying TEs differentially expressed shared between the scions grafted onto *S. torvum* (TOR) and tomato ‘Emperador RZ’ (EMP), data in parenthesis indicate the total number of differentially expressed TEs in each condition. **c** LTR identity values of *de novo* annotated intact LTR-TEs commonly regulated in both hetero-grafted scions, separated in up-regulated (UP) and down-regulated (DOWN) elements. The red lines represent averaged values.

To further investigate LTR TEs expression, we developed the functional annotation pipeline *LTRpred* (https://hajkd.github.io/LTRpred/) and applied it to the eggplant genome assembly to *de novo* re-annotate LTR TEs. We designed *LTRpred* to screen for old and young LTR TEs and to predict their functional capacity based on a well-defined sequence composition and intact sequence motifs. Together, *LTRpred* allowed us to study the association between novel LTR TEs and their epigenetic regulation during grafting. Differential expression analysis of these newly annotated LTR TEs again correlated in both hetero-grafted combinations (**Fig. S4c-d**, **S7**), similarly to what we previously observed for genes (**Fig. 3**). A down-regulation trend was observed for the annotated LTR TEs both in plants grafted onto *S. torvum* (73%) and ‘Emperador RZ’ (66%) compared to the self-grafted controls (**Table S11-S12**). Specifically, in eggplant scions grafted onto *S. torvum*, 63 differentially expressed LTRs were identified (17 are up-regulated and 46 down-regulated) (**Fig. S4c-d**), while 32 TEs were differentially expressed in plants grafted onto ‘Emperador RZ’ rootstock (11 up-regulated and 21 down-regulated) (**Fig. S4c-d**). A significant proportion of these LTR TEs (6 up-regulated genes and 20 down-regulated) were shared between the two heterograft combinations (**Fig. S4c-d, Table S11-S12)**.

Interestingly, we observed that the average LTR identity, a strong indicator of the age of TEs, is high (= young elements) in LTR-TEs upregulated in hetero-grafted scions and consistently lower (= old elements) in down-regulated LTR-TEs (**Fig. 4c**). Therefore, our result suggests that the heterograft condition might regulate the transcription of TEs in an age-dependent fashion, by promoting expression of older TEs and by repressing young and potentially more mobile TEs.

## Discussion

Eggplant is one of the most successful commercially grafted herbaceous plants, with a high degree of compatibility for interspecific grafting which may provide enhanced vigour and resistance to pathogens (Lee and Oda, 2010). While most resistance to root pathogens of grafted plants derives from intrinsic properties of the rootstocks, the molecular mechanism of grafting and how particular graft combinations enhance scion vigour is largely unknown.

Grafting experiments in the model plant *Arabidopsis* have revealed that during long-distance movements, transgene-derived and endogenous sRNAs move across a graft union and are able to direct DNA methylation in the genome of the recipient cells, inducing physiological changes (Melnyk, 2017; Molnar et al., 2010). Here, using eggplant as model for grafting-induced vigour, we found that genome-wide CHH hypomethylation in the scions correlates with enhanced plant vigour. Although we could not identify direct effects of DNA methylation changes on gene expression, two hetero-grafted scions displayed a similar expression profile, characterized by up-regulation of genes involved in cell division and down-regulation of genes involved in secondary metabolism and defence. Interestingly, a similar transcriptional change pattern is reported in hybrids for many heterotic plant species, and it is generally associated to an increase in plant vigour (Blum, 2013). In the last decade, there has been a growing appreciation of the potential role of epigenetics in the molecular, cellular, and developmental bases of heterotic vigour (Catoni and Cortijo, 2018; Groszmann et al., 2011). In previous studies performed in *Arabidopsis*, heterotic vigour was observed in the hybrid progeny obtained by crossing near-isogenic parents with variable epigenetic profiles (Dapp et al., 2015; Lauss et al., 2018), indicating that the genetic difference in the parents is not the only factor triggering heterosis. In our study, we observed that a methylation decrease in CHH context was associated to vigour in hetero-grafted eggplant scions, suggesting that changes in DNA methylation induced by the rootstocks in the scion can contribute to an increase in vigour. Remarkably, a genome-wide decrease in CHH methylation was previously associated to hybrid vigour in *Arabidopsis*, and correlates with general decrease of 24 nt siRNAs (Greaves et al., 2012; Groszmann et al., 2011).

In plants, methylation in CHH context is normally associated with suppression of TEs expression. Therefore, it is surprising that the observed genome-wide decrease of methylation in hetero-grafted scions does not correlate with a wide increase of TE expression, but is rather associated with a more complex regulation resulting in many TEs being down-regulated. One possible explanation is that hypomethylation might activate other silencing mechanisms to reduce RNA transcripts of potentially active TEs, for example Post Transcriptional Gene Silencing (PTGS). This hypothesis is consistent with the observed up-regulation of AGO7, associated to PTGS in *Arabidopsis* (Carbonell and Carrington, 2015), and the preferential suppression of younger and potentially more active LTR TEs.

Our work provide the first DNA methylome of eggplant, and shed light to the molecular mechanisms underlying the effect of rootstock on scion. Our data suggest the involvement of epigenetic regulation to control vigour of grafted Solanaceae species, showing that epigenetic changes (especially decrease in CHH methylation) correlate with vigour in two hetero-grafted eggplant combinations, mirroring the well-known effects reported for hybrids and epi-hybrids. In this context, the use of grafting represents a promising alternative to traditional breeding to manipulate plant epigenomes and improve plant production.

## Materials and Methods

### Plant material and sampling

Eggplant double haploid (DH) line derived from the commercial hybrid ‘Ecavi’ (Rijk Zwaan, Netherlands), wild *Solanum torvum* and tomato F1 *S. lycopersivum* × *S. habrochaites* hybrid ‘Emperador RZ’ (Rijk Zwaan, Netherlands) (Bogoescu and Doltu, 2015) plants have been selected for this work. ‘Ecavi’ plants were used as self-grafted controls and as scions for the following rootstock/scion grafting combinations: ‘Ecavi’/’Ecavi’ (self-grafted control), ‘Emperador RZ’*/* ‘Ecavi’, *S. torvum/* ‘Ecavi’. Seeds were sterilized as described by Gisbert *et al*., 2006 and germinated in growth chambers under long-day conditions (26°C, 16-h light, 8-h dark) (Gisbert et al., 2011). Grafting was performed using the cleft methods (Gisbert et al., 2011) and moved in an experimental greenhouse of Carmagnola, Italy (44°53′N; 7°41′E) 3 months after grafting, during the 2017 growing season. Scion leaves of 3 biological replicates for each grafted and control plant type were sampled 5 months after grafting, flash-frozen in liquid nitrogen and stored at −80 °C.

### Phenotypic evaluation

Hetero-grafted and control plants have been phenotypically monitored every week in the experimental green house. Plant height and scion developmental architecture were annotated, and the sampling time was selected as the moment of largest vigour difference between hetero-grafted and self-grafted combinations.

### Nucleic Acid Extraction

DNA and RNA were extracted from 100 mg of frozen leaves tissue collected from eggplant scions or ungrafted plants. For each sample, genomic DNA was extracted using the Qiagen Plant DNeasy kit (Qiagen, Hilden, Germany). Total RNA was extracted using the Spectrum Plant Total RNA Kit (Sigma, Saint Louis, USA) method according to the manufacturer’s instructions.

### Bisulfite conversion of genomic DNA

Genomic DNA (120 ng) were bisulfite-converted using the EZ DNA Methylation-Gold Kit (Zymo Research, Irvine, CA) following the manufacturer’s recommendation with minor modifications. In order to increase the chances to obtain a high conversion rate, the conversion step was repeated twice. Samples underwent the following reaction in a thermal cycler: 98°C for 10 minutes, 64°C for 2.5 hours, 98°C for 10 minutes, 53°C for 30 minutes, then 8 cycles at 53°C for 6 minutes followed by 37°C for 30 minutes and a final incubation at 4°C overnight. Bisulfite conversion was performed on duplicates of each experimental condition.

### Library preparation and sequencing

Converted samples were immediately used to prepare bisulfite libraries employing the TrueSeq DNA Methylation Kit (Illumina, San Diego, CA) accordingly to the protocol’s instruction. Libraries for RNA expression analysis were prepared in duplicates from 2 μg of total RNA using the TrueSeq Stranded mRNA Sample Prep Kit (Illumina, San Diego, CA) following the manufacturer’s instructions. Libraries quality and fragment sizes were checked with a TapeStation 2200 (Agilent technologies, Santa Clara, CA) instrument and the DNA quantified by PCR on a LightCycler 480 II (Roche Molecular Systems, Pleasanton, CA) using the Library Quantification Kit (Roche Molecular Systems, Pleasanton, CA). Bisulfite and RNA sequencing reactions were performed on a NextSeq500 using a HighOutput chemistry, at the core facility of the Sainsbury Laboratory University of Cambridge (SLCU, Cambridge, UK).

### Sequencing processing

Whole genome bisulfite sequencing (WGBS) and RNA-seq raw reads were trimmed using Trimmomatic (Bolger et al., 2014) to remove adapter sequences. For bisulfite libraries, high-quality trimmed sequences (on average 90,7 % of raw reads) were aligned against the eggplant reference genome (Barchi et al., 2019) using Bismark (Krueger and Andrews, 2011). Genome coverage was estimated taking into consideration the following parameters: length of reads, read numbers and eggplant genome size (~1.2 Gb). Not repeated DNA regions of the eggplant chloroplast sequence (Ding et al., 2016) were used to estimate the bisulfite conversion. We computed chloroplast mappability on the eggplant genome using the gem-mappability tool from the Gem library (Derrien et al., 2012), with a k-mer size of 75 bp and allowing a maximum of one mismatch, and only unique regions (mappability = 1) were used to estimate conversion. To account for non-converted DNA, we applied a correction according to Catoni et al. (2017). Briefly, the number of methylated reads were decreased as: m*= max(0, m – nc) (where m* is the corrected number of methylated reads, m is the raw number of methylated reads, n is the total number of reads and c is the conversion rate). DNA methylation at different cytosine contexts were plotted on chromosome using the R package DMRcaller (Catoni et al., 2018).

For transcript level analysis, reads were mapped with TopHat (Trapnell et al., 2009) on the eggplant reference genome (Barchi et al., 2019), using parameters–max-multihits 1–read-realign-edit-dist 0–no-mixed. Mapped reads were subsequently counted using htseq-count (Anders et al., 2015) with parameters–order name–type = exon–stranded = reverse. As most of plant genomes are composed of transposable elements and LTRs are the most abundant, we developed the functional annotation pipeline *LTRpred* (https://hajkd.github.io/LTRpred/) to *de novo* re-annotate retrotransposons within the eggplant genome assembly. The output of *LTRpred* was then used as input for htseq-count. We applied a stringent presence-call filter, restricting the analysis to those annotated genes or LTRs with more than five counts-per-million in the two biological replicates. Differential expression was assessed with DEseq (Anders and Huber, 2010), using as thresholds log2 fold change > 1 and a Benjamini-Hochberg’s FDR < 0.05.

### GO enrichment analysis

We performed a GO enrichment analysis using ShinyGO online tool (Ge and Jung, 2018) with Fisher’s exact test, false discovery rate (FDR) correction and selecting a 0.05 p-value cut-off. This approach was employed for the analysis of statistically significant DE genes and LTR-TEs.

### Target expression analysis

RNA-seq data were validated using qPCRs for 11 genes up or down regulated according to FPKM values. For real-time qRT-PCR analysis, total RNA (2 μg) was treated with RQ1 DNase (Promega, Madison, Wisconsin) and reverse-transcribed with the SuperScript VILO cDNA Synthesis Kit (Thermo Fisher, Waltham, Massachusetts), according to the manufacturer’s instructions. PCRs were carried out in duplicate using 10 ng of template cDNA, 10 nM target-specific primers (**Table S5**) and LightCycler 480 SYBR Green I Master (Roche Molecular Systems, Pleasanton, CA) in the LightCycler 480 II detection system (Roche Molecular Systems, Pleasanton, CA) in a volume of 10 μl. GADPH was used as housekeeping gene.

## Data availability

Sequencing data have been deposited in Gene Expression Omnibus under the accession number (data under submission process).

## Author contribution

S.L. conceived the research. M.C., C.C. and E.C. designed the experiments. E.C. performed experiments. J.P. and C.G. planned and handled the grafting of the plants. D.V. managed the plant growth in field, E.P. performed phenotypical data collection and analysis. E.C., M.C., L.B. and H.G.D. analysed data. E.C. and M.C. wrote the manuscript with contributions from C.C. S.L., J.P., H.G.D, C.G. E.P revised the manuscript. All authors read and approved the final manuscript.

## Acknowledgements

Part of the computations described in this paper were performed using the University of Birmingham’s Compute and Storage for Life Sciences (CaStLeS) service. This work was supported by European Research Council (EVOBREED) [322621]; Gatsby Fellowship [AT3273/GLE]; University of Birmingham personal starting grant of M. Catoni [GBGB BIP1267].

We are grateful to Dr. J. Paszkowski (Sainsbury Laboratory, Cambridge, UK) and all the members of his research group for the support and fruitful scientific discussions during the experimental work, Dr. J. Griffiths (Sainsbury Laboratory, Cambridge, UK) for the critical reading of the manuscript and R. Schina (FMI, Basel, Switzerland) for helping in the first stages of eggplant development and for software assistance.

---

**Figure S1.**
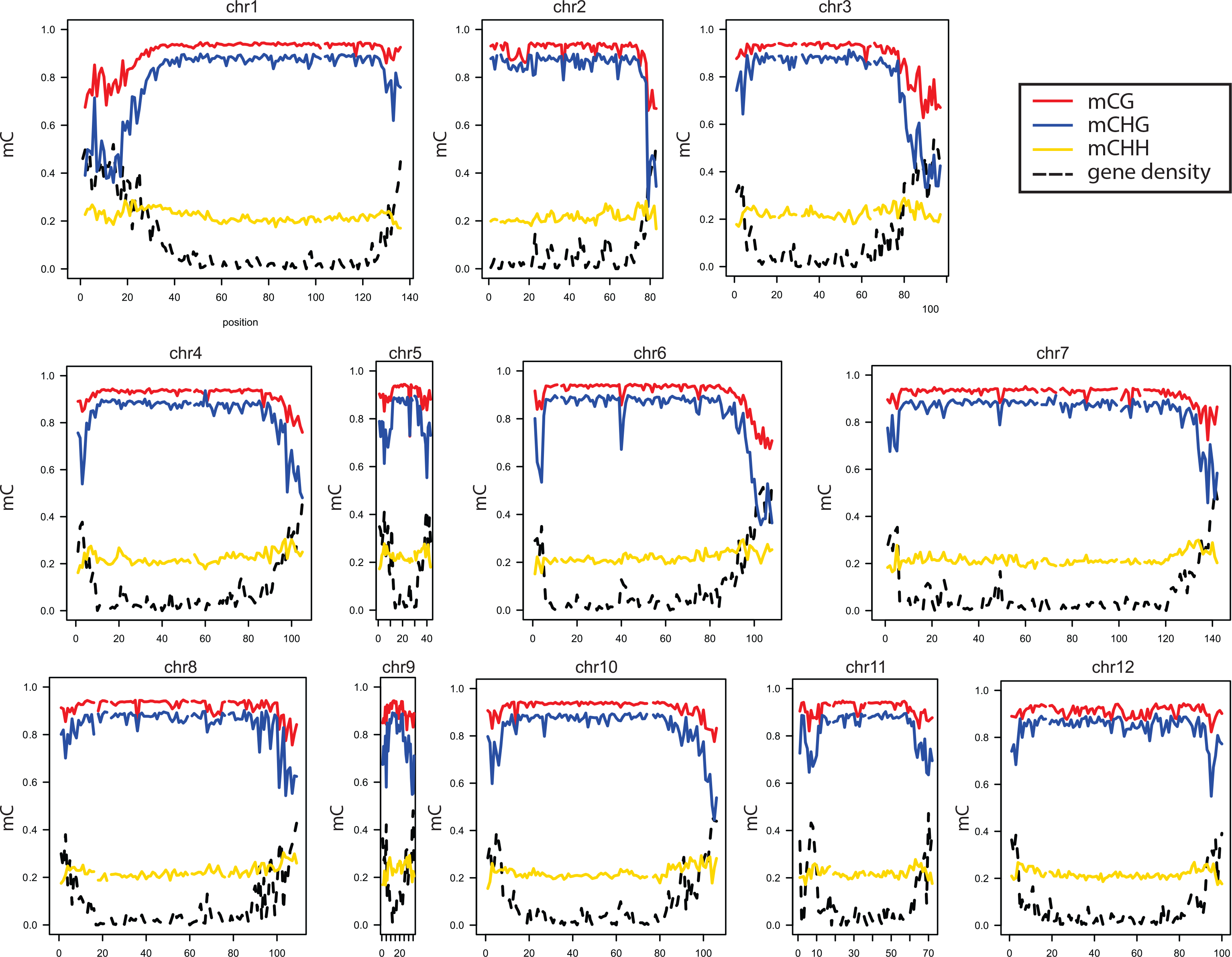
DNA methylation profile in eggplant leaves of not grafted plants. DNA methylation content in CG (red), CHG (blue) and CHH (yellow) contexts, represented in the 12 eggplant chromosomes. In dashed black is reported the gene density.

**Figure S2.**
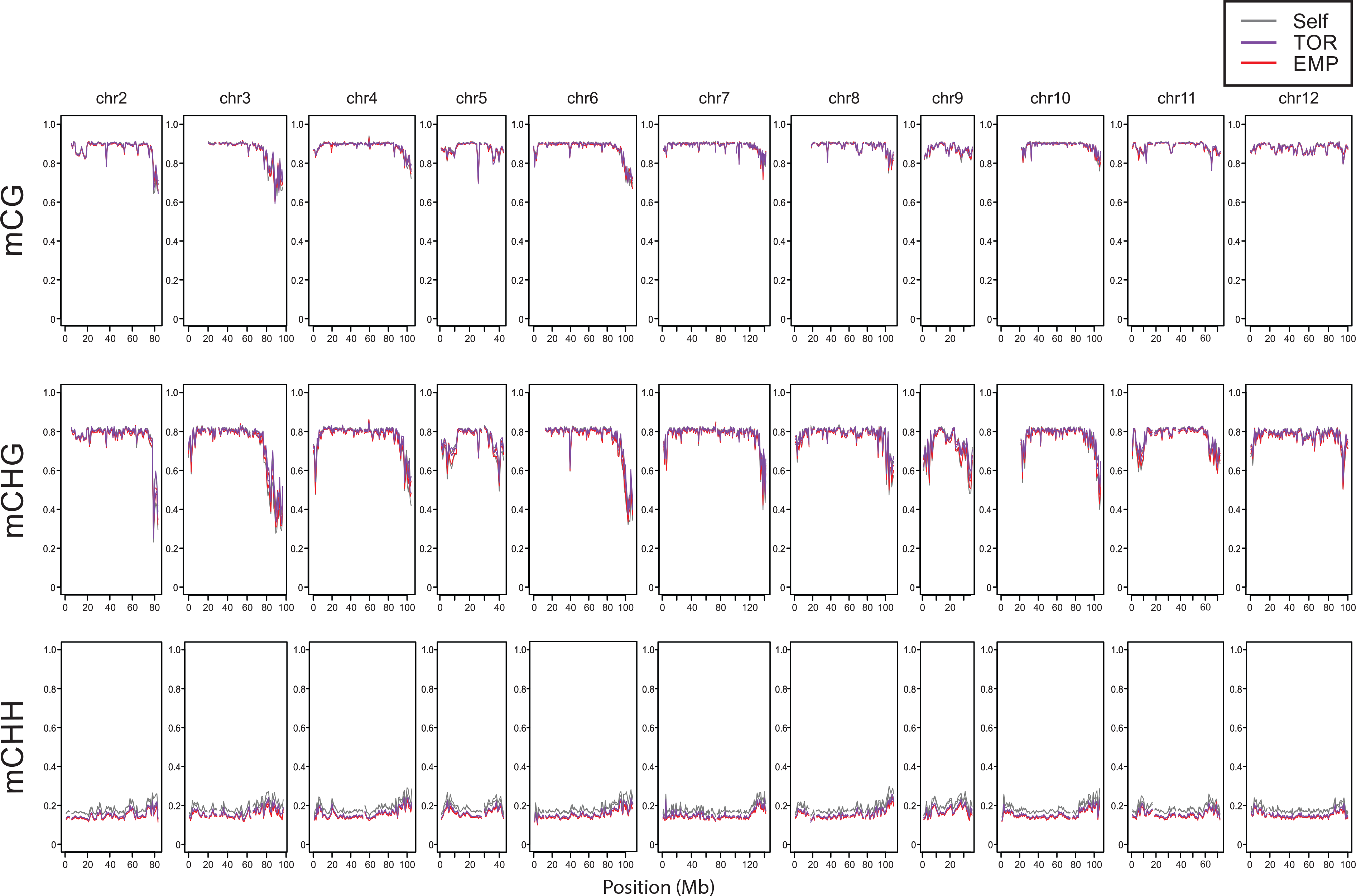
DNA methylation profiles in self-grafted and hetero-grafted eggplant. Distribution of DNA methylation at each cytosine context (mCG, mCHG and mCHH) along chromosomes 2 to 12, in eggplant scions self-grafted (Self), or grafted onto *S. torvum* (TOR) or tomato ‘Emperador RZ’ (EMP) rootstocks. The profiles in chromosome 1 are displayed in **Figure 2**.

**Figure S3.**
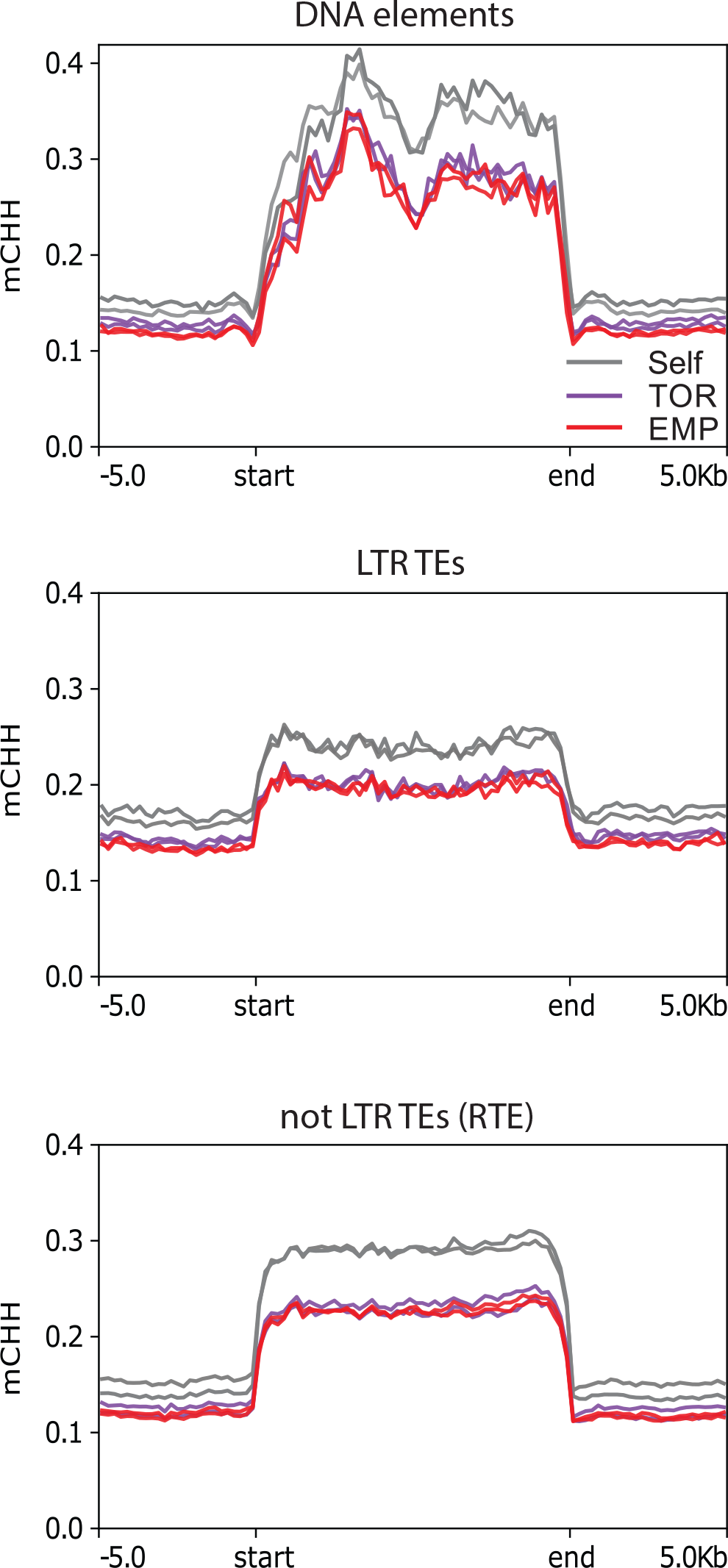
DNA methylation distribution at the main TEs classes. Distribution of averaged DNA methylation (at CG, CHG and CHH contexts) at annotated TEs of main three groups in eggplant genome, in eggplant scions self-grafted (Self), grafted on *S. torvum* (TOR) or tomato ‘Emperador RZ’ (EMP).

**Figure S4.**
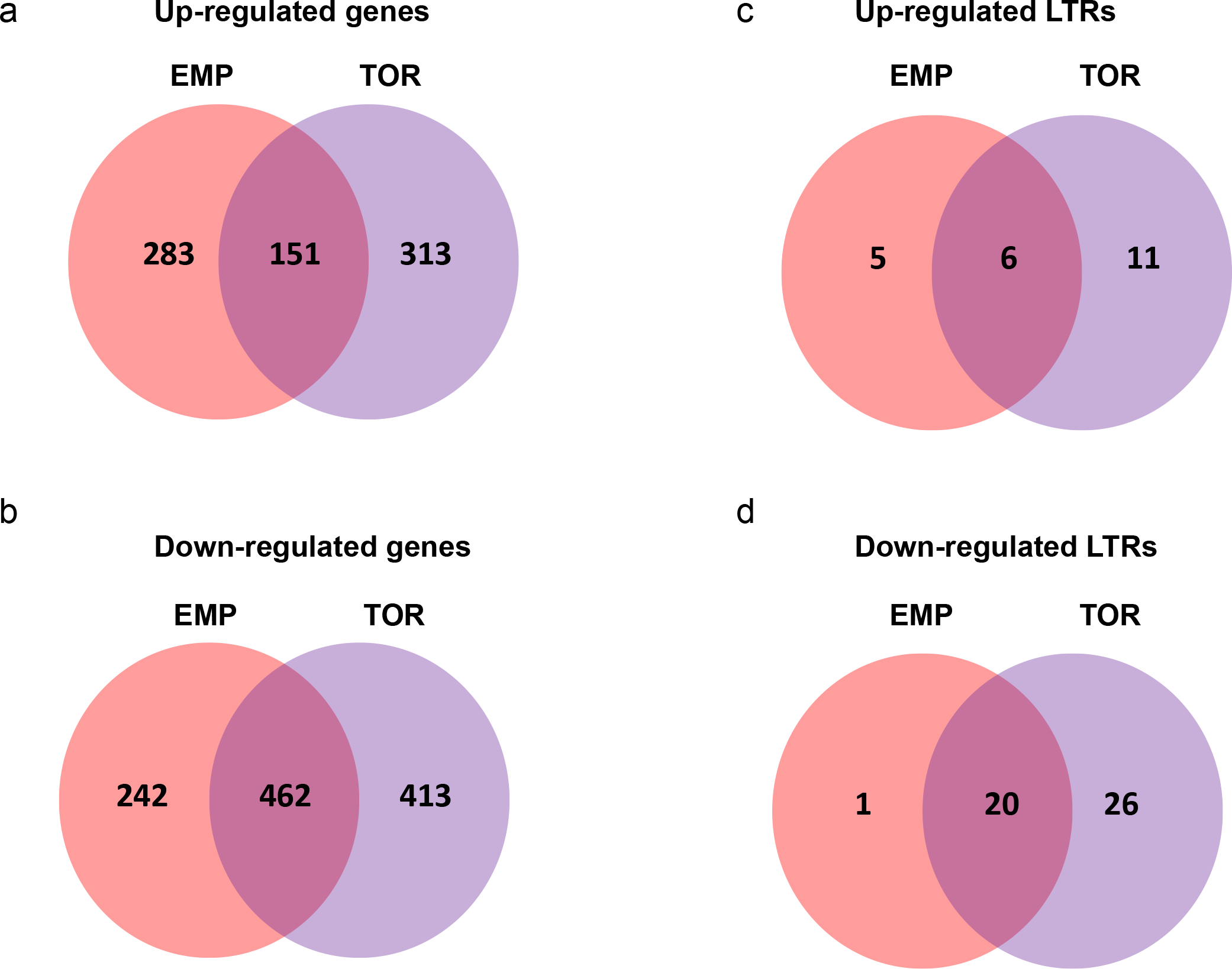
Distribution of differentially expressed annotated genes and LTR-TEs. Venn-diagrams displaying up-regulated (**a**) and down-regulated (**b**) genes, or up-regulated (**c**) and down-regulated (**d**) annotated LTR-TEs, in eggplant scions grafted onto *S. torvum* (TOR) and tomato ‘Emperador RZ’ (EMP) compared to self-grafted control.

**Figure S5.**
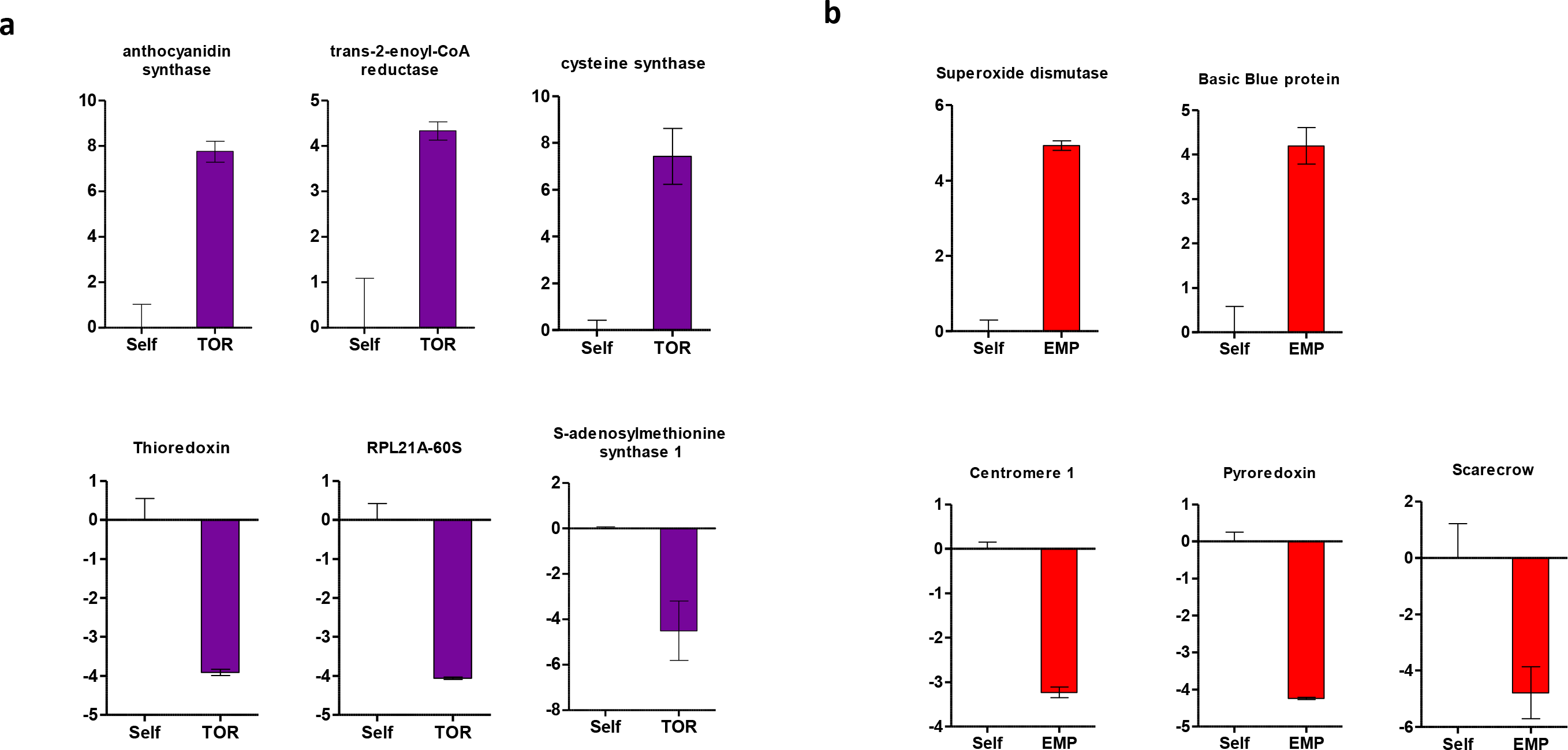
Validation of RNA-seq expression data. RNA accumulation was measured with qPCR for a subset of 11 genes randomly selected with |log2FC| > 3. Bars represent up/down-regulated genes in eggplant scions grafted onto *S. torvum* (**a**) or tomato ‘Emperador RZ’ (**b**) rootstocks compared to self-grafted plants (Self). Both up-regulated (top line) and down-regulated (bottom line) genes were tested. Error bars represent SD of three replicates.

**Figure S6.**
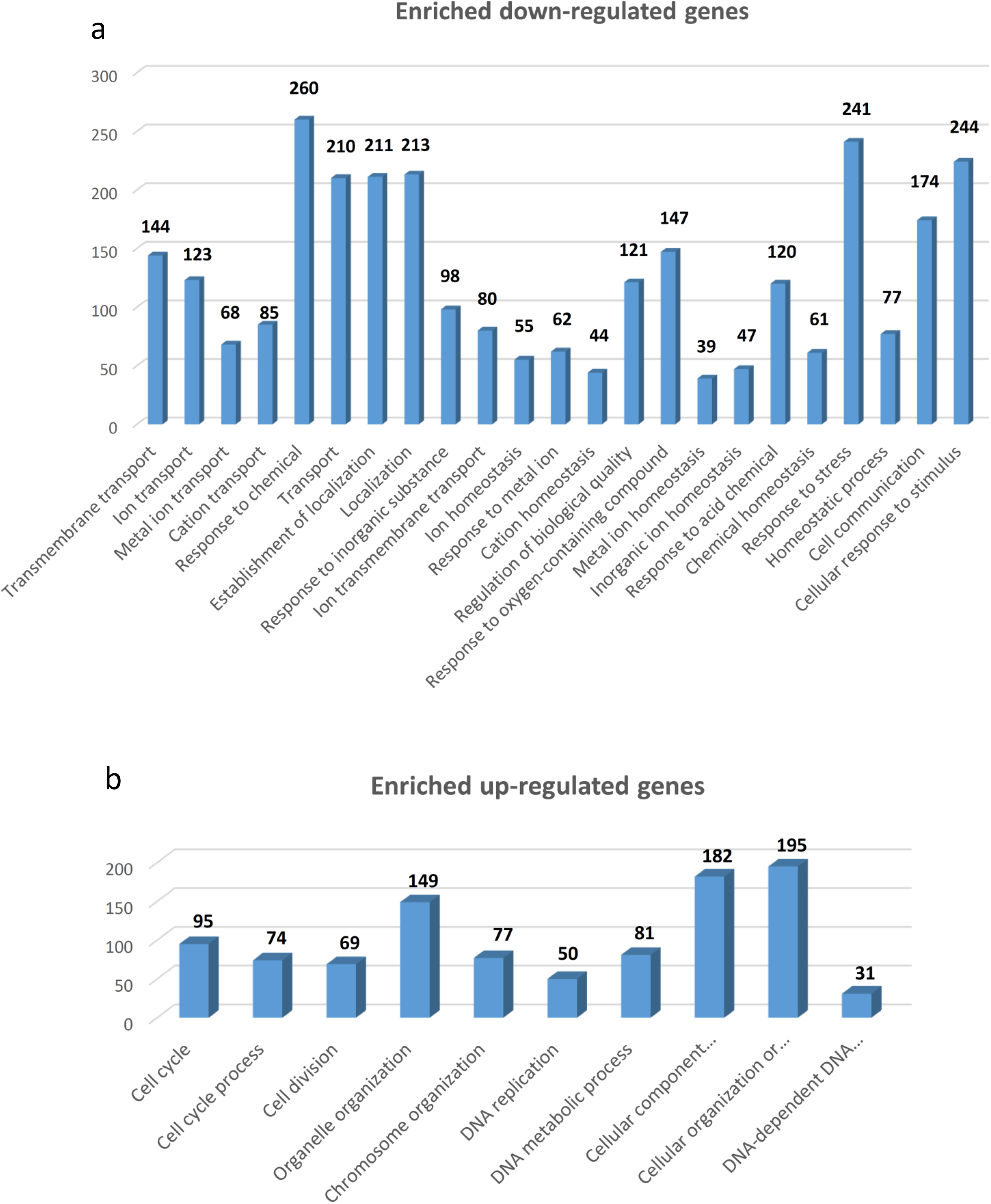
Enrichment genes analysis in self-grafted and hetero-grafted eggplants. Lists of enriched Gene ontology categories of differentially expressed genes shared in heterograft compared to self-grafted scions. Different graph display information for down-regulated (**a**) and up-regulated (**b**) genes. The number of genes belonging to each category is reported at the top of each bar.

**Figure S7.**
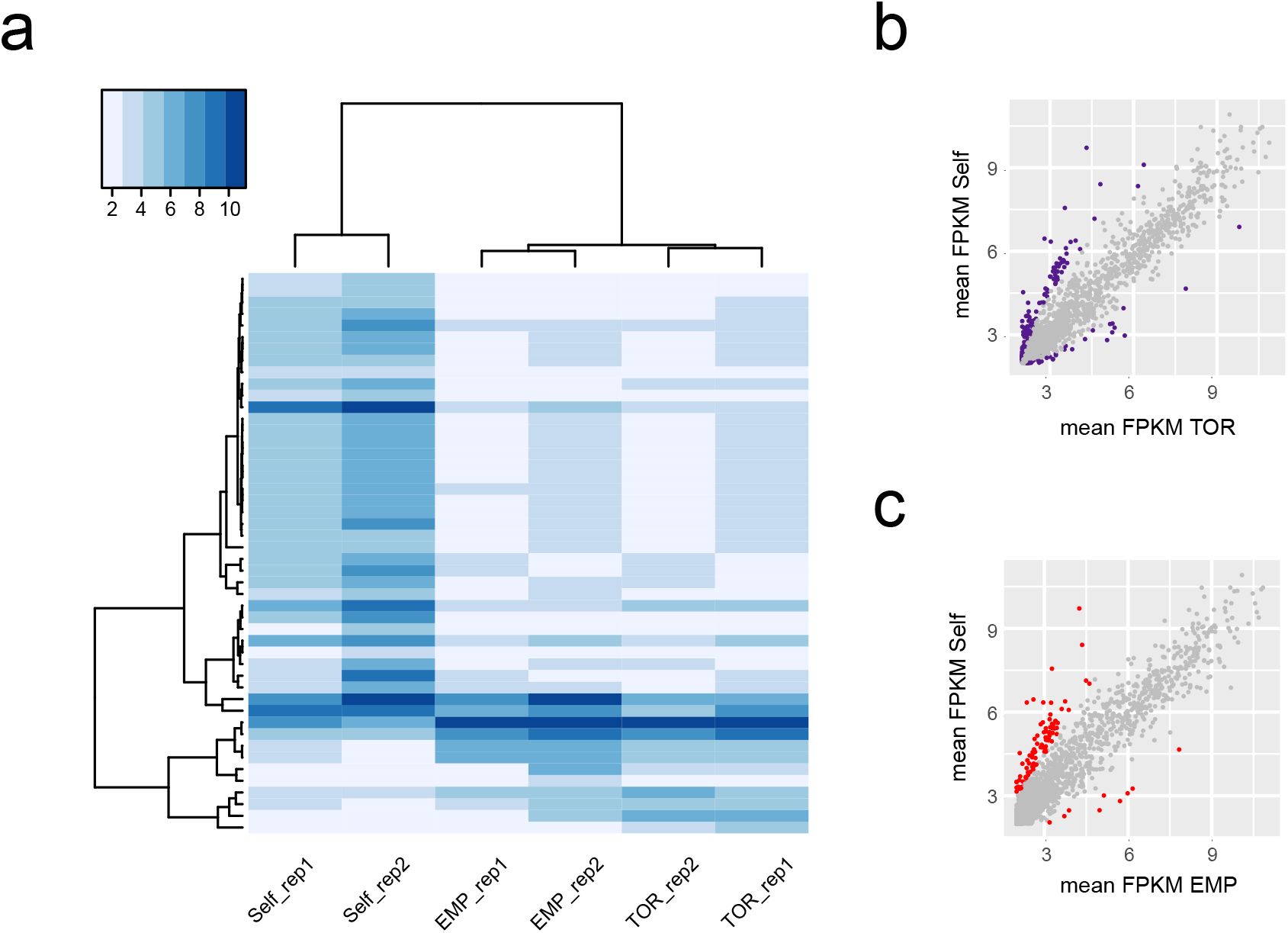
Hetero-grafted scions display similar LTR-TEs expression profiles. **a** Heatmap illustrating the expression (log2 FPKM) of differentially expressed *de novo*-annotated LTR TEs (|log2| > 2 and FDR <0.05) in hetero-grafted plants and self-grafted controls. Each row represents one LTR TE (*n* = 63). Coloured bars on the top-left indicate the expression level. Each column represents an eggplant scion grafted on different rootstock (TOR = *S. torvum*, EMP = tomato ‘Emperador RZ’ and Self = self-grafted). Expression values were ordered according to hierarchical clustering (hclust and heatmap3 R software environment). The Euclidean distance dendrogram is presented on the left. **b** Scatter-plots of annotated LTR-TEs (*n* =6,583) in Self and EMP conditions, each dot represents a genes and in red are displayed differentially expressed LTR-TEs between the two conditions. **c** Scatter-plots of annotated LTR-TEs (*n* =6,583) in Self and TOR conditions. In purple are displayed differentially expressed LTR-TEs between the two conditions.

